# *Saccharomyces cerevisiae* strain YH166: a novel wild yeast for the production of tropical fruit sensory attributes in fermented beverages

**DOI:** 10.1101/149732

**Authors:** Kara Osburn, Robert Caputo, Justin Miller, Matthew L. Bochman

## Abstract

All ales are fermented by various strains of *Saccharomyces cerevisiae.* However, recent whole-genome sequencing has revealed that most commercially available ale yeasts are highly related and represent a small fraction of the genetic diversity found among *S. cerevisiae* isolates as a whole. This lack of diversity limits the phenotypic variations between these strains, which translates into a limited number of sensory compounds created during fermentation. Here, we characterized a collection of wild *S. cerevisiae*, *S. kudriavzevii*, and *S. paradoxus* strains for their ability to ferment wort into beer. Although many isolates performed well, *S. cerevisiae* strain YH166 was the most promising, displaying excellent fermentation kinetics and attenuation, as well as a tropical fruit sensory profile. Use of this strain in multiple styles of beer suggested that it is broadly applicable in the brewing industry. Thus, YH166 is a novel ale strain that can be used to lend fruity esters to beer and should pair well with citrusy hops in hop-forward ales.

## 1. Introduction

Many fermented beverages, including beer, wine, kombucha, and kefir, are produced (at least in part) by the metabolic action of yeasts in the genus *Saccharomyces* (1). These organisms are ubiquitous in such applications due to their naturally high levels of tolerance to ethanol (EtOH), low pH, osmotic stress, and anaerobic conditions (2-4). Of the eight species in the genus *Saccharomyces* (*S. arboricola*, *S. cerevisiae*, *S. eubayanus*, *S. jurei*, *S. kudriavzevii*, *S. mikatae*, *S. paradoxus*, and *S. uvarum*) (5), *S. cerevisiae* is most commonly found in traditionally fermented beverages and is used industrially for beverage fermentation and bioethanol production (1,6). Indeed, *S. cerevisiae* is used for ale and wine production worldwide.

However, recent research has shown that the *S. cerevisiae* strains currently used for these processes lack in genetic variability, with the *Saccharomyces pastorianus* strains used for lager production displaying even less diversity. Three different groups performed whole genome sequencing of 212 (7), 157 (8), and 28 (9) *S. cerevisiae* strains, most of which are used to make beer and wine. These studies found that industrial strains are polyphyletic, representing multiple (in the case of ale strains, three (8,9)) domestication events. However, among the individual clades of *S. cerevisiae* strains representing German, British, and wheat beer isolates, the yeasts are extremely genetically similar and display high levels of inbreeding (7). Further, they represent only a small fraction of the natural genetic diversity found among other sequenced examples of *S. cerevisiae*. This narrow genotypic and phenotypic variation among commercially available strains likely limits the spectrum of sensory compounds produced by these yeasts during fermentation. Notably, in beers that are not heavily hopped, the yeast can account for ≥ 50% of the sensory profile of the finished beverage (10).

As such, much work has be performed to engineer strains with improved or altered fermentation and sensory compound production profiles. For instance, strains have been engineered to reduce haze development caused by polyphenolic compounds in beer (11) and eliminate the off-flavor compound diacetyl (12). However, many of these manipulations generate yeasts that are considered genetically modified organisms (GMOs) because selectable markers and heterologous genes are introduced into the *S. cerevisiae* genome. As there is a public bias against consuming GMO products (13), more recent efforts have focused on breeding and hybridization methods to produce yeasts that are viewed as non-GMOs. Indeed, several groups have made great strides in this area (14-16). Such projects tend to be labor- and cost-intensive though. An alternate to constructing new brewing strains with desired sensory attributes is bio-prospecting for wild isolates with these phenotypes. Indeed, wild *S. cerevisiae* strains represent an untapped reservoir of aromas and flavors in beverage fermentation. We define wild strains as those isolated via open fermentation (*i.e.*, air capture) or from environmental samples (*e.g.*, soil and plant matter).

Here, we characterized a collection of wild *Saccharomyces* strains (17,18) for their evolutionary relatedness to each other and commercially available ale strains, their ability to metabolize mono- and disaccharides, their EtOH tolerance, and ability to ferment wort into palatable beer. Of the isolates tested, *S. cerevisiae* strain YH166 stood out for its excellent fermentation kinetics, EtOH and osmotic stress tolerance, and pleasing sensory attributes that were reminiscent of tropical fruit. We suggest that the use of wild strains such as YH166 for beverage fermentation will represent the next trend in the ongoing global craft beverage revolution.

## 2. Materials and methods

### 2.1. Strains, media, and other reagents

*S. cerevisiae* strains WLP001 and WLP300 were purchased from White Labs (San Diego, CA). Wild strains were isolated as described in (17). All yeast strains were grown on yeast extract, peptone, and dextrose (YPD; 1% (w/v) yeast extract, 2% (w/v) peptone, and 2% (w/v) glucose) plates containing 2% (w/v) agar at 30°C and in YPD liquid culture at 30°C with aeration unless otherwise noted. All strains were stored as 15% (v/v) glycerol stocks at -80°C. Media components were from Fisher Scientific (Pittsburgh, PA, USA) and DOT Scientific (Burnton, MI, USA). All other reagents were of the highest grade commercially available.

### 2.2. Phylogenetic analysis

The wild *Saccharomyces* strains were identified at the species level by sequencing the variable D1/D2 portion of the eukaryotic 26S rDNA as described (18). After species identification, the phylogenetic relationships among the strains were determined by aligning their rDNA sequences using ClustalX (19). The alignments were iterated at each step but otherwise utilized default parameters. ClustalX was also used to draw and bootstrap neighbor-joining (N-J) phylogenetic trees using 1000 bootstrap trials; the trees were visualized with TreeView v. 1.6.6 software (http://taxonomy.zoology.gla.ac.uk/rod/rod.html). The *Schizosaccharomyces pombe* rDNA sequence (GenBank accession HE964968) was included in the alignments as the outgroup, and this was used to root the N-J tree in TreeView. WLP001 and WLP300 were included to determine the relatedness of the wild strains to commercially available ale yeasts. Sequences for other *Saccharomyces* species were retrieved from the Nucleotide database of the National Center for Biotechnology Information (NCBI; https://www.ncbi.nlm.nih.gov/nucleotide/).

### 2.3. Sugar metabolism

The yeast strains were grown by inoculating 5 mL YPD liquid medium with single colonies from YPD plates and incubation overnight at 30°C with aeration. The optical density at 660 nm (OD_660_) of each culture was determined using a Beckman Coulter DU730 UV/Vis Spectrophotometer. Then, the cells were diluted to an OD_660_ = 0.1 in 200 μL YPD medium containing 2% (w/v) glucose, maltose, sucrose, or xylose in round-bottom 96-well plates, overlaid with 50 μL mineral oil to prevent evaporation, and incubated at 30°C with shaking in a BioTek Synergy H1 plate reader. The OD_660_ of every well was measured and recorded every 15 min for 14-15 h, and these values were plotted *vs.* time to generate growth curves. All growth experiments were repeated ≥ 3 times, and the plotted points represent the average OD_660_ values. Error bars representing standard deviations were omitted for clarity.

### 2.4. EtOH and glucose tolerance

Ethanol tolerance was measured as above but in 96-well plates containing YPD liquid medium or YPD liquid medium supplemented with 5, 10, or 15% EtOH. Glucose tolerance was likewise assessed in 96-well plates containing YPD liquid medium (2% (w/v) glucose) or YP liquid medium supplemented with 10, 20, or 30% (w/v) glucose.

### 2.5. Test fermentations

Laboratory-scale fermentations were performed as described (18). Briefly, the yeast strains were grown to saturation in 4 mL of YPD liquid medium and used to inoculate ~400 mL of blonde ale wort (1.050 original gravity (OG)) in 500-mL glass fermenters fitted with standard plastic airlocks. The fermenting cultures were incubated at ~22°C for 2 weeks. Un-inoculated wort was treated as above to control for wort sterility. Prior to bottling into standard 12-oz brown glass bottles, their final gravity (FG) was measured using a MISCO digital refractometer (Solon, OH). Flocculation was qualitatively judged by comparing the clarity of the experimental beers to control beers fermented with *S. cerevisiae* WLP001 or WLP300, which routinely display medium or low flocculation, respectively (http://www.whitelabs.com/yeast-bank). Bottle conditioning was conducted as in (20) at room temperature for ≥ 2 weeks. The comparisons between WLP001 and YH166 fermentations were conducted in 1-L glass cylinders (30 cm tall, 7.5 cm inner diameter) for 6-7 days at an average temperature of 23.6 ± 0.3°C. The gravity and alcohol by volume (ABV) were monitored in real time using BeerBug digital hydrometers (Sensor Share, Richmond, VA). Small-batch (15-20 L) fermentations were performed by the Bloomington Hop Jockeys (http://hopjockeys.org) home brewing club (Bloomington, IN). Multiple worts were produced for these trials, which are detailed in the Supplementary Materials.

## 3. Results

### 3.1. *Phylogeny of the wild* Saccharomyces *strains*

As we previously reported (17,18), we have isolated hundreds of wild yeasts with the potential for use in the beverage fermentation industry (Supplemental Table 1). Among the strains isolated, many *Saccharomyces* species were uncovered, including 37 *S. cerevisiae*, eight *S. paradoxus*, and one *S. kudriavzevii* (18). Because some of these samples came from locations in and around production breweries, we screened them for potential contamination by commercial strains of brewer’s yeast. To do so, we aligned the D1/D2 region of their rDNA and constructed a phylogenetic tree to compare the evolutionary relatedness of the isolated strains with commercial controls and previously annotated D1/D2 sequences from multiple *Saccharomyces* species (Fig. 1). We found that the yeasts clustered into five distinct clades (I-V), with the wild *S. paradoxus* and *S. kudriavzevii* strains all contained within Clade IV. The two commercial strains WLP001 and WLP300 were grouped into Clade V and appear to be closely related to the YH196 and WYP75 isolates, respectively. This suggests that YH196 and WYP75 could be commercial contaminants, and thus, they were excluded from further analyses. It is unclear why the *S. cerevisiae* isolates clustered into four clades (I-III and V) rather than a single group.

**Figure 1.**
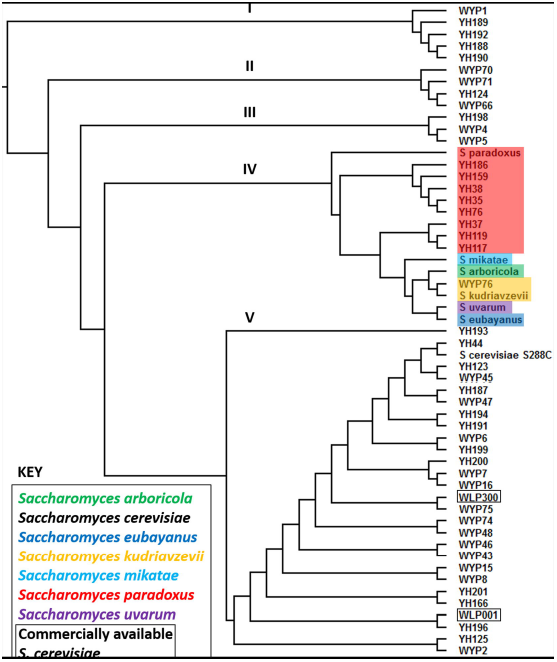
Evolutionary relationships among the wild *Saccharomyces* strains and two commercially available ale yeasts. The D1/D2 rDNA sequences of the indicated strains were aligned, and the phylogenetic relationships among them were drawn as a rooted N-J tree using *Schizosaccharomyces pombe* (ATCC16979) as the outgroup. Five distinct clades of strains are marked with Roman numerals. The *S. paradoxus* strains are highlighted red, *S. mikatae* is light blue, *S. arboricola* is green, the *S. kudriavzevii* strains are orange, *S. uvarum* is purple, and *S. eubayanus* is dark blue. All *S. cerevisiae* strains are unhighlighted, though the two commercial controls (WLP001 and WLP300) are boxed. Sequence data for strains without YH or WYP designations were retrieved from the NCBI with the following accession numbers: *S. paradoxus* KT972121, *S. mikatae* AB040996, *S. arboricola* JQ914741, *S. kudriavzevii* AB040995, *S. uvarum* KJ469964, *S. eubayanus* LT594193.1, and *S. cerevisiae* S288C NR_132207.1.

### 3.2. Characterization of sugar metabolism, ethanol tolerance, and flocculation.

To begin to triage the isolated *Saccharomyces* strains for those most likely to perform well in beer fermentation, we sought to characterize their abilities to metabolize various sugars, their ethanol tolerance, and how well they flocculate. First, to assay for sugar metabolism, we followed the growth of each strain in rich medium (YP) supplemented with 2% (w/v) of two common monosaccharides (glucose and xylose) and disaccharides (maltose and sucrose). We found that the wild isolates could be phenotypically categorized into four groups (Supplemental Table 1), representatives of which are shown in Figure 2A-D. Yeasts in Group 1 could equivalently utilize the preferred sugars glucose and sucrose but displayed little-to-no growth in the presence of xylose and maltose (Fig. 2A). Strains in Group 2 were likewise able to metabolize glucose and sucrose, as well as displayed an intermediate level of growth in medium containing maltose (Fig. 2B). The isolates in Group 3 displayed similar growth kinetics and cell densities in the presence of glucose, maltose, and sucrose but weak growth in xylose-containing medium (Fig. 2C). Finally, the yeasts in Group 4 grew well in the presence of all four tested carbon sources but achieved the highest cell densities in medium containing glucose or sucrose (Fig. 2D). It should be noted that *S. cerevisiae* is generally considered incapable of metabolizing xylose, but it does encode an endogenous xylose utilization pathway that can be activated by the over-expression of the nonspecific aldose reductase *GRE3* and the xylitol dehydrogenase *XYL2* genes (21). We hypothesize that *GRE3* and *XYL2* are naturally upregulated in the Group 3 and 4 yeasts that grew in the presence of xylose as the sole carbon source.

**Figure 2.**
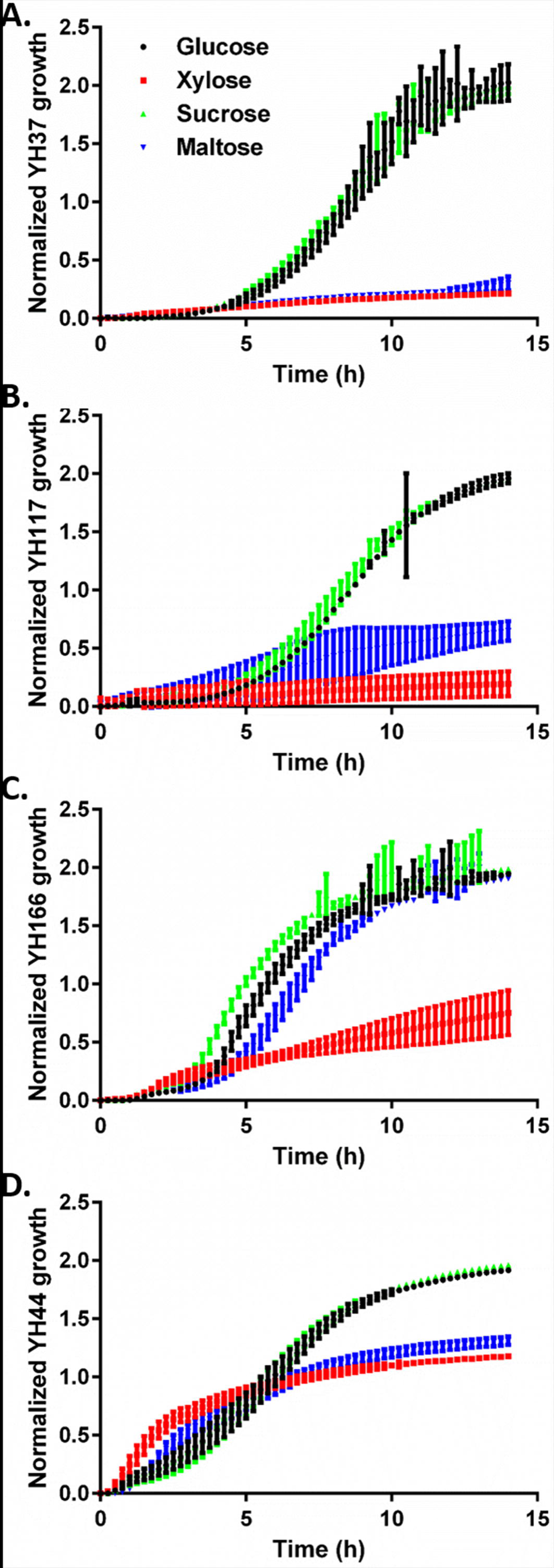
Growth curves of representative strains utilizing various sugars. Small liquid cultures of the strains indicated on the y-axes were grown in 96-well plates in YP medium supplemented with 2% (w/v) of the indicated sugars. The OD_660_ of each well was monitored with a plate reader. Four different phenotypes were found: A) yeasts in Group 1 grew poorly in the presence of xylose and maltose; B) yeasts in Group 2 displayed a moderate level of growth in the presence of maltose; C) yeasts in Group 3 grew very well in the presence of maltose; and D) yeasts in Group 4 grew well in the presence of all tested sugars. The plotted points in each curve represent the average OD_660_ values of ≥ 3 independent experiments normalized to the highest OD_660_ in each experiment. The error bars are the standard deviation.

Next, ethanol tolerance was similarly assessed by growing strains in YPD medium containing 0-15% ABV. Again, the various strains could be grouped based on their growth curves. As shown in Figure 3A, some strains were insensitive to increasing EtOH concentrations, growing as rapidly and to nearly as great a density in the presence of 15% ABV as in the complete absence of EtOH. Other strains displayed similar sensitivities to all concentrations of EtOH tested, though growth was still evident (Fig. 3B). However, most strains displayed a concentration-dependent sensitivity to EtOH, with higher ABVs increasingly inhibiting growth (Fig. 3C). Regardless, all strains grew to some extent in the presence of 15% ABV (Fig. 3 and data not shown), corresponding to the well-documented natural EtOH tolerance of *Saccharomyces* species (2-4).

**Figure 3.**
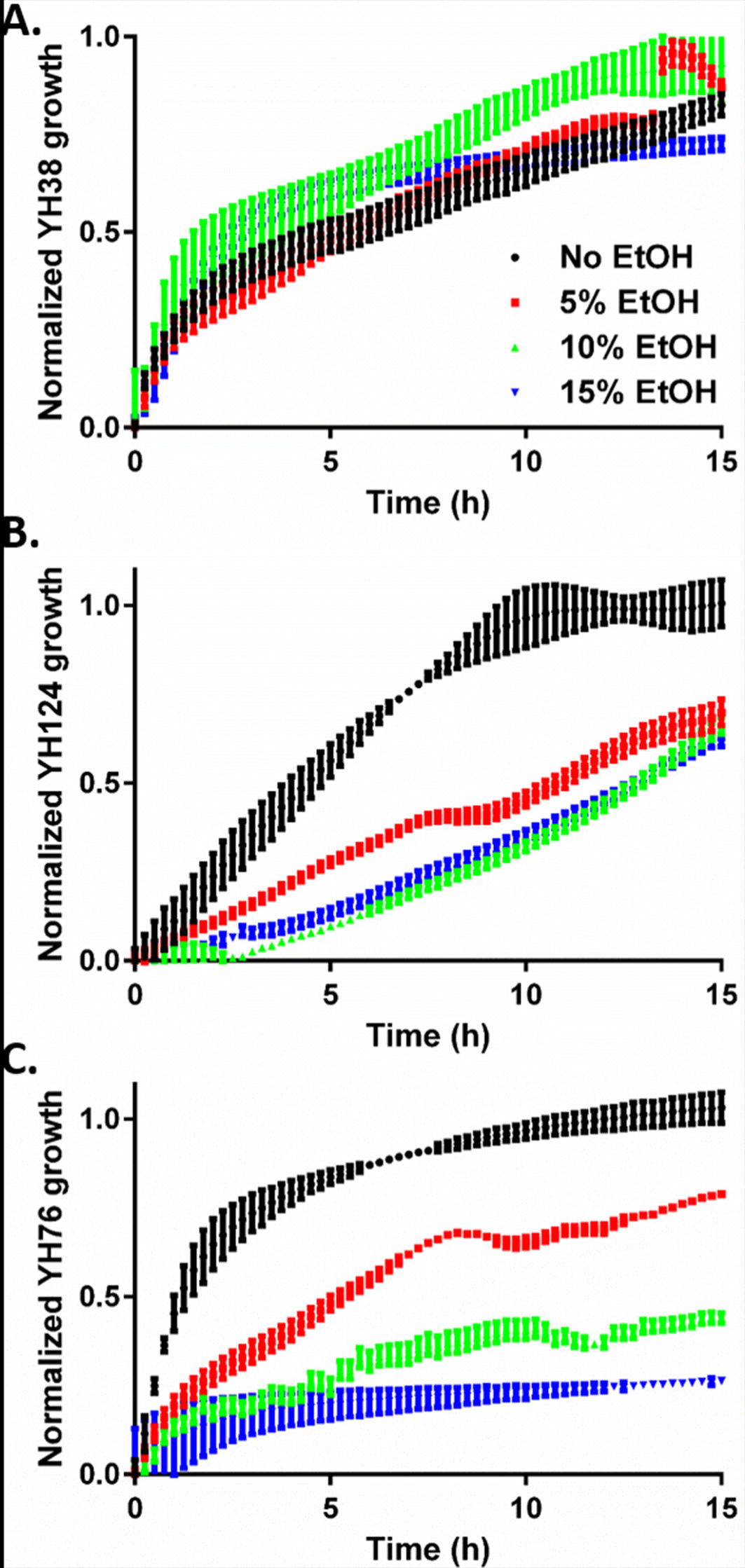
EtOH tolerance of representative strains. Small liquid cultures of the strains indicated on the y-axes were grown in 96-well plates in YPD medium or YPD medium supplemented with the indicated amount of EtOH. The OD_660_ of each well was monitored with a plate reader. Four different phenotypes were found: A) tolerance of EtOH up to 15% (v/v), B) similar sensitivities to 5-15% EtOH, and C) concentration-dependent sensitivity to EtOH. The plotted points in each curve represent the average OD_660_ values of ≥ 3 independent experiments, and the error bars are the standard deviation.

Flocculation was qualitatively assessed by comparing the rate of cell sedimentation by the wild strains to two commercial controls WLP001 and WLP300 (medium and low flocculation, respectively, see www.whitelabs.com) in small stationary liquid cultures and in small fermenters. In both cases, all of the wild strains displayed medium or higher levels of flocculation (Table 1, Supplementary Table 1, and data not shown). However, we did also note that some of the strains formed rather loose slurries that were easily disrupted, sending cells back into suspension with only gentle agitation.

**Table 1.** Laboratory-scale fermentation results of select strains.

| Strain | Species<sup>a</sup> | Attenuation<sup>b</sup> | Flocculation | Sensory |
| --- | --- | --- | --- | --- |
| YH35 | *S. paradoxus* | 27% | High | Chlorophenol/bandage |
| YH37 | *S. paradoxus* | 44% | Medium | Chlorophenol/bandage |
| YH38 | *S. paradoxus* | 41% | Medium | Chlorophenol/bandage, vegetal |
| YH44 | *S. cerevisiae* | 44% | Medium | Clean, neutral |
| YH76 | *S. paradoxus* | 41% | Medium | Chlorophenol/bandage |
| YH124 | *S. cerevisiae* | 31% | Medium | Neutral |
| YH166 | *S. cerevisiae* | 70-80% | High | Green apple, guava, dry, slightly tart, tropical fruit |
| WYP2 | *S. cerevisiae* | 65-80% | Medium | Clean, neutral, reminiscent of lager |
| WYP4 | *S. cerevisiae* | 61% | Medium | Silky mouthfeel, cereal grain notes |
| WYP5 | *S. cerevisiae* | 61% | Medium | Saison-like |
| WYP6 | *S. cerevisiae* | 75-85% | Medium | Thin, farmhouse-like, pithy, earthy |
| WYP7 | *S. cerevisiae* | 70-80% | Medium | Crisp, dry, slightly bitter, reminiscent of champagne |
| WYP43 | *S. cerevisiae* | 60-80% | Medium | Clean, fruity esters, Belgian phenolic |
| WYP76 | *S. kudriavzevii* | 68% | Medium | Neutral |
| WLP001 | *S. cerevisiae* | 70-80% | Medium | Clean, neutral, slightly fruity |

<sup>a</sup> All species are in the *Saccharomyces* genus.
<sup>b</sup> Strains displaying a range of attenuation were brewed with up to six times to delineate the range. The others were brewed with at least two times, and the greatest attenuation is reported.

### 3.3. Small-scale fermentations

Aside from the strains that metabolized maltose poorly (*e.g.*, YH37; Fig. 2A), the other wild isolates all displayed good beer fermentation potential based on our initial tests. To begin to characterize the brewing capacity of these strains, we performed small wort fermentations with each. We utilized WLP001 as a positive control for levels of attenuation and flocculation, as well as a baseline for our sensory analyses. After two rounds of test brewing and analysis, we chose the most promising strains for additional trials. The full data set can be found in Supplementary Table 1, and representative strains are shown in Table 1.

We found that the *S. paradoxus* isolates ranged in their ability to attenuate from 20-55% (Table 1 and Supplementary Table 1) with an average attenuation across all strains of ~37%. Aside from under-attenuation, the beers produced by *S. paradoxus* all smelled and tasted heavily of adhesive bandages, which was likely due to the production of chlorophenol (22). Thankfully, only two *S. cerevisiae* strains (WYP15 and WYP16) shared this sensory phenotype (Supplementary Table 1). Overall, the *S. cerevisiae* strains displayed better attenuation (average of 69%), though they varied widely from 17-95%. Many of the beers produced were neutral in aroma and flavor, though some were fruity, had a Belgian strain phenolic character, and/or were slightly tart and reminiscent of saison or farmhouse ales. The single isolate of *S. kudriavzevii* attenuated well (68%) and yielded neutral sensory characteristics (Table 1).

Of all of the wild *Saccharomyces* strains that we tested, YH166 repeatedly displayed good brewing characteristics, with excellent attenuation (70-80%), flocculation, and aroma/flavor production (Table 1). In every tasting panel that we conducted, the sensory profiles of the beers made by YH166 were consistently characterized as “tropical”, with notes of guava and green apple. Other strains also displayed similar attenuation and flocculation, but the beers they produced were generally neutral in sensory and comparatively bland when sampled alongside beer fermented by YH166. Thus, we focused on YH166 for further characterization.

### 3.4. Brewing with YH166

*S. cerevisiae* YH166 was isolated from a spontaneous fermentation conducted in a vacant lot during the summer of 2015 in Indianapolis, IN (18). This wild fermented beer contained six distinct yeast strains: three isolates of *Brettanomyces bruxellensis* and one strain each of *Candida zeylanoides*, *S. cerevisiae* (YH166), and *Wickerhamomyces anomalus*. YH166 was the fastest growing and most vigorous fermenting strain of the six under laboratory conditions (data not shown). Indeed, when compared to WLP001 in laboratory-scale fermentations, YH166 reliably reached terminal attenuation >24 h faster, though its terminal ABV (~5.5%) was always slightly less than that produced by WLP001 (~6%; Fig. 4A).

**Figure 4.**
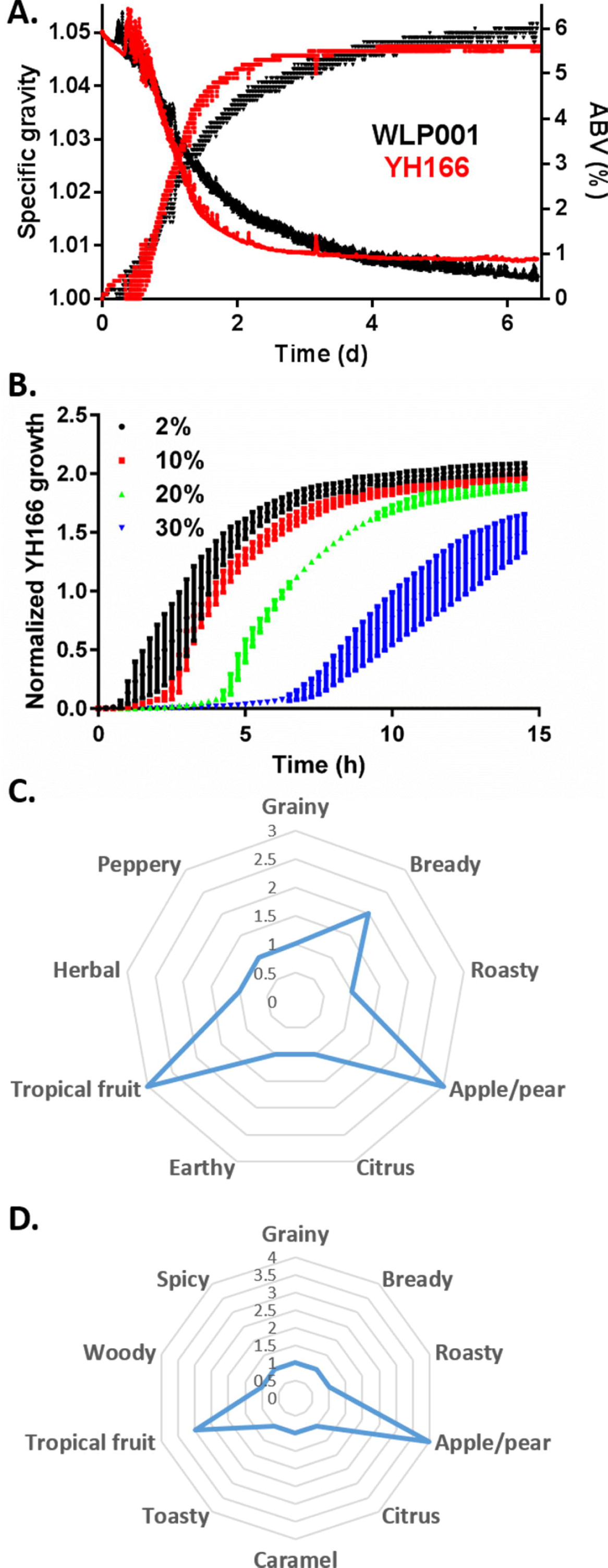
YH166 displays rapid fermentation kinetics and tolerance to osmotic stress. A) *S. cerevisiae* strains YH166 and WLP001 were independently inoculated into fermenters containing a 1.050 OG wort, and the fermentation kinetics were followed in real time using Wi-Fi-enabled digital hydrometers. SG is plotted on the left y-axis and ABV (%) on the right. The data shown are representative of three independent fermentations for each strain. B) Small liquid cultures of YH166 were grown in 96-well plates in YP medium supplemented with the indicated concentrations of glucose. The OD_660_ of each well was monitored with a plate reader. The plotted points in each curve represent the average normalized OD_660_ values of ≥ 3 independent experiments, and the error bars are the standard deviation. C) Spider plot of aroma descriptors for beers fermented with YH166 (see Supplementary Materials for details). D) Spider plot of flavor descriptors for beers fermented with YH166 (see Supplementary Materials for details).

We typically use low gravity wort for laboratory-scale fermentations. However, ale strains are commonly utilized in a variety of beer styles, some of which have very high OGs, such as Russian imperial stouts (23). To determine if YH166 could tolerate high gravity wort, we assessed the growth of this strain in rich medium containing 2-30% (w/v) glucose. As shown in Figure 4B, the lag time to exponential growth increased from < 1 h in 2% glucose to > 5 h in 30% glucose. However, YH166 was able to overcome the osmotic stress of the glucose at all concentrations and grow to high density, suggesting that it is suitable for fermenting worts with a wide range of OGs.

Finally, we assessed the activity of YH166 in a variety of worts and fermentation conditions (see Supplementary Materials) with the help of the Bloomington Hop Jockeys, a local home brewing club. It should be noted that each fermentation experiment was only performed a single time, but we feel that the range of conditions tested is still worthy of report. Consistent with our laboratory-scale fermentations, YH166 performed well in all of these trials and produced aromatic (Fig. 4C) and flavor profiles (Fig. 4D) that were reminiscent of apple/pear and tropical fruit. Contrary to the laboratory-scale experiments, however, these beers were uniformly cloudy or hazy in appearance (data not shown).

## 4. Discussion

The natural tolerance displayed by *Saccharomyces* species to fermentation stresses such as ethanol, low pH, and anaerobic growth (2-4) have enabled these organisms to dominate most industries that rely on fermentation worldwide. However, many of the yeast strains that are currently used in these processes are highly genetically related (7,8). We sought to characterize wild *Saccharomyces* strains for their ability to ferment wort into beer to determine if novel sensory characteristics can be found in the untapped array of yeast isolates present in nature.

Based on phylogenetics (Fig. 1) and phenotypic analyses (Fig. 2-3, Tables 1, and Supplementary Table 1), the strains in our collection of wild yeasts could be divided into a variety of groups. It was our hope that one or more of the phylogenetic groups would be indicative of isolates with positive fermentation attributes to help direct future yeast hunting efforts. This largely proved not to be the case though. For instance, phylogenetic clade IV was dominated by *S. paradoxus* strains that fermented poorly and/or produced unpalatable beer (Fig. 1), but clade IV also contained *S. kudriavzevii* WYP76, which produced quaffable beer. The only strain grouping that was relevant for beer fermentation was Group 1 in sugar metabolism (Fig. 2A). Yeasts in Group 1 utilized maltose poorly and consequently attenuated poorly during fermentation (Table 1 and data not shown). Such isolates will be avoided during our ongoing yeast bio-prospecting by only selecting for strains that can rapidly metabolize maltose.

Our current results also suggest that *S. paradoxus* strains should be avoided for beer fermentation. All eight tested here created a repulsive aroma and taste that was reminiscent of adhesive bandages (Table 1). This is a common off-flavor in beer production that is attributable to chlorinated phenols (24). Very little has been reported in the scientific literature about brewing with *S. paradoxus*, and it has been suggested that this is one of the only *Saccharomyces* species not used commercially for fermentation (25). Perhaps this dearth of information is due to off-putting sensory profiles produced by *S. paradoxus* strains. A brief survey of online resources indicated that home brewers and craft brewers have successfully used *S. paradoxus* in brewing without encountering an antiseptic or medicinal sensory profile, but these reports cannot be verified. Regardless, we collected all of our *S. paradoxus* strains from the bark of oak trees (18), so chlorophenol production appears to be a common characteristic of wild *S. paradoxus* isolated from this natural reservoir.

Unlike the *S. paradoxus* strains, most of the remaining *Saccharomyces* isolates tested produced beers with neutral or more flavorful and pleasing sensory profiles (Table 1). Not all of them attenuated to high levels, but flocculation matched or exceeded the WLP001 control. Serial re-inoculation of low-attenuating strains into wort for fermentation may help to “domesticate” such strains by adapting them to beer production (26), and ongoing experiments are investigating this issue. Many strains, such as YH166, were well suited to fermentation with no manipulation other than the process of enrichment and pure culturing (17).

We chose to focus on strain YH166 due to its excellent fermentation kinetics and tropical fruit sensory profile. In our laboratory-scale trials, it performed as well as the WLP001 ale control strain (Fig. 4A) and demonstrated excellent resistance to osmotic stress (Fig. 4B), suggesting that it can be used to ferment beers with high OGs. YH166 was also amenable to a variety of beer styles when used by home brewers (see Supplementary Materials) and consistently produced sensory profiles with apple/pear and tropical fruit notes. Interestingly, the home brew experiments uniformly yielded beers that were hazy or cloudy in appearance, in contrast to the high flocculation we found in the laboratory (Table 1). Many factors affect flocculation (reviewed in (27)), and thus additional experiments should be performed to determine the effects of variables such as pH, wort gravity, temperature, and cations on YH166 flocculation. Regardless, this lack of flocculation coupled with otherwise desirable brewing characteristics and fruity sensory attributes suggests that YH166 may be an attractive strain for New England-style India pale ale (NE-IPA) brewing. Indeed, NE-IPAs are cloudy-to-opaque and generally described as juicy and fruity (28). Thus, novel wild brewing strains such as YH166 can make an immediate impact on current trending styles of beer and could lead to the development of new beer styles based around the yeast as the core ingredient.

## Acknowledgements

We thank the members of the Bloomington Hop Jockeys home brewing club for providing valuable brewing data for strain YH166, as well as members of the Bochman laboratory for critically reading this manuscript and providing feedback. This work was supported by startup funds from Indiana University, a Translational Research Pilot Grant from the Johnson Center for Entrepreneurship in Biotechnology, a grant from the American Homebrewers Association Research and Education Fund, and funds from Wild Pitch Yeast, LLC (to MLB).

